# Escaping through the predator’s gill cleft: A defensive tactic of juvenile eels after capture by predatory fish

**DOI:** 10.1101/2021.06.08.447630

**Authors:** Yuha Hasegawa, Kazuki Yokouchi, Yuuki Kawabata

## Abstract

Some prey animals can actively escape from predators even after being captured, but knowledge of such behaviors is still limited, especially in vertebrates. Here, we report the unique active escape behavior of Japanese eel *Anguilla japonica* juveniles through the gill cleft of the predatory fish *Odontobutis obscura*. Of the 54 *A. japonica* juveniles captured by the predator, 28 escaped via the predator’s gill, and most escaped individuals survived afterwards. In all escaped individuals, their tails emerged first from the gill cleft of the predator, and then their whole bodies slipped out in a backward direction from the cleft. These findings indicate that *A. japonica* juveniles have the specialized antipredator tactic, which provides the basis for further investigation from behavioral, ecological, and evolutionary perspectives.

---

Predation is a strong selective force that shapes various forms of defensive tactics of prey. Most previous studies have focused on the defensive tactics of prey before being captured by predators (e.g., counter-adaptations against searching, recognition, and catching by predators) [1], and “capture” is often considered a synonym for death of prey. Nonetheless, some animals exhibit passive defensive tactics after being captured, such as the use of spines, toxins, and safe passage through the digestive system [1], and recent studies have shown that specific invertebrates can actively escape from the digestive systems of predators [2, 3]. Here, we report the unique active escape behavior of Japanese eel *Anguilla japonica* juveniles after being captured by the predatory fish *Odontobutis obscura*, involving escape through the predator’s gill cleft.

Because of their high commercial value, unique morphology (e.g., elongated body form), and ecology (e.g., long-distance migration), a large number of studies have been conducted on temperate anguillids (e.g., Japanese eels *A. japonica*, European eels *A. anguilla*, and American eels *A. rostrata*) [4]. However, surprisingly little is known about their defensive tactics against predators. For fishes with a pelagic larval phase, the predation mortality at the post-settlement juvenile stage is typically selective (rather than random) and is highly dependent on antipredator behavioral traits such as escape response [5]. Therefore, we initiated a predator–prey interaction experiment using *A. japonica* juveniles and the sympatric piscivorous predator *O. obscura* in an aquarium setting.

In the preliminary experiment, we discovered a unique defensive behavior of *A. japonica* juveniles after being captured by the predator, where they escaped through the predator’s gill cleft. To verify that this behavior is an active defensive tactic rather than a rare, anecdotal observation, we proceeded predator–prey interaction experiment and obtained detailed information regarding the escape success rate, escape sequence, and time required to escape (see Supplemental Information).

In general, the predator attacked *A. japonica* by opening its mouth, and the predator moved toward the *A. japonica* individual while the *A. japonica* was drawn toward the predator’s mouth owing to suction. Of the 54 *A. japonica* individuals the predator had captured, 28 individuals (51.9%) escaped via the predator’s gill 47.4 ± 36.2 s after capture (Fig. 1; Video S1–S3; Table S1). In all the escaped cases, the individual’s tails emerged first through the gill cleft of the predator regardless of the body part (head, center, or tail) initially attacked, and then their whole bodies slipped out in a backward direction from the cleft. There were no significant effects of initially attacked body parts on escape success or the time to escape (generalized linear mixed model, p > 0.1). When the tail of *A. japonica* emerged from the predator’s gill cleft, the predator often showed some resistance behaviors (e.g., swimming around the aquarium, and rubbing the body against the aquarium wall; Video S2). Most escaped individuals (26 out of 28 individuals) survived for at least 48 h. Severe scratch wounds were observed in one individual after the escape, which was subsequently euthanized by an overdose of 2-phenoxyethanol. No apparent wounds were observed in the other individuals, but one individual died during the observation period.

**Fig. 1.**
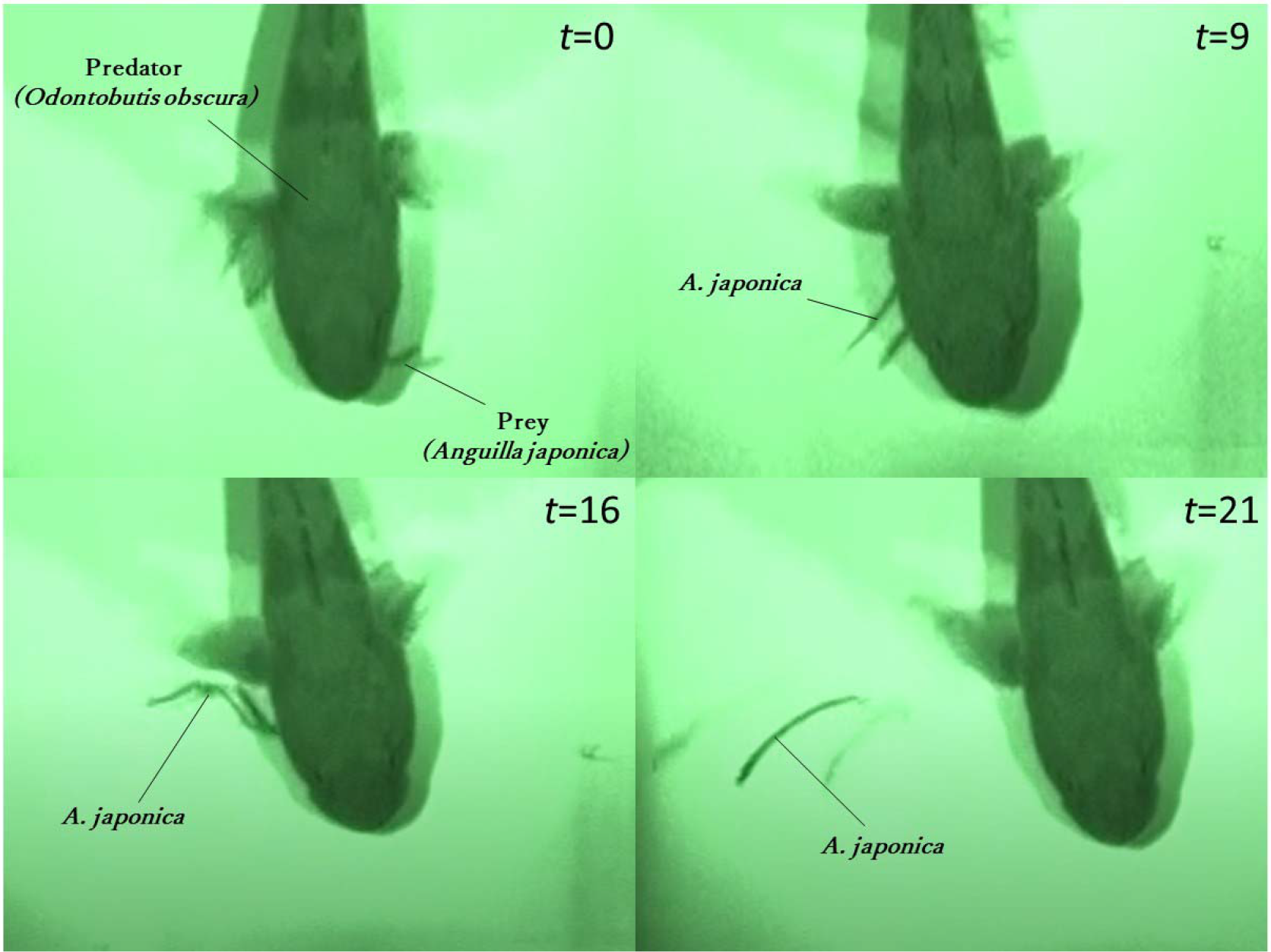
Representative behavioral sequence of Japanese eel *Anguilla japonica* juveniles after being captured by the predatory fish *Odontobutis obscura*, recorded using an infrared video camera. After the predator captured the *A. japonica* individual (*t* = 0 s), the tail of *A. japonica* first emerged through the gill cleft of the predator (*t* = 9 s), the whole body of *A. japonica* gradually slipped out from the cleft (*t* = 16 s), and finally the *A. japonica* individual swam away from the predator (*t* = 21 s). Note that the shadows of *A. japonica* and *O. obscura* are reflected on the bottom of the tank.

Interestingly, all escaped *A. japonica* individuals escaped leading with their tail parts, regardless of their initially attacked body parts. This behavior could be related to the unique locomotion style of this species. Unlike the escape response of many fish species (i.e., C-start), anguilliform species quickly retract their heads to evade sudden threatening stimuli [6]. Additionally, the congeneric species *A. anguilla* uses backward swimming as a burst escape response in dark and complex habitats [7]. Therefore, it is possible that *A. japonica* has a fixed behavioral trait (i.e., its tail is inserted into the predator’s gill cleft, and then the fish moves backward to escape through it) to facilitate escape from the predator via the gills. Another possibility is that some of the captured individuals had tried to escape headfirst and were not successful, but we could not observe the behavioral pattern because it occurred in the buccal cavity of the predator. Further research on *A. japonica* behaviors in the predator’s buccal cavity (such as by X-ray videography) is needed to clarify the behavioral and kinematic features of the escape behavior via the predator’s gill.

The elongated body form has evolved convergently in many aquatic animals [6], but its underlying selective force (i.e., adaptive significance) is still unclear. Our results suggest that natural selection would favor an elongated body form in cases where prey must escape through gaps in the predator’s organs, as prey with deeper/wider bodies would not be able to pass through these gaps. Consistent with this idea, the parasitic gordian worm, which has an extremely elongated body, can avoid predation by escaping via the mouth, nose, and gill of the predator that had captured its insect host [3]. It would be interesting to test whether other elongate species (both phylogenetically close and distant species to *A. japonica*) possess defensive tactics similar to those observed in *A. japonica*. This will provide insights into the evolution of the elongated body form in aquatic animals.

After the oceanic pelagic phase, *A. japonica* and other temperate eels settle in rivers, estuaries, and coastal seas, where they encounter predators [4, 8, 9]. Although our knowledge of predation on *A. japonica* in its early life stage is limited, channel catfish *Ictalurus punctatus* and blackfin sea bass *Lateolabrax latus* were found to prey on *A. japonica* during its settlement phase [9]. These predators use suction feeding to engulf whole prey into their mouths [10], similar to that by *O. obscura*. Thus, the observed *A. japonica* defensive tactic would be effective for escaping from these predators as well, at least to some extent. In addition to predatory fish with suction-feeding style, birds and water rats were found to prey on the juveniles of other eel species [8]. Further research to determine eel predators and predation experiments using different predator types are required to clarify the extent to which this defensive tactic contributes to the survival of juvenile eels in the wild.

Our findings indicate that *A. japonica* juveniles have a specialized antipredator tactic, and prey–predator battles occur in the predator’s buccal cavity. Many prey species possess antipredator tactics at the phases of encounter, detection, and attack but not after capture. Thus, this unique behavior will provide an additional opportunity to avoid predation for *A. japonica* and possibly other elongate species.

## Supplemental information

Supplemental Information includes one table, three videos, experimental procedures, acknowledgments, author contributions, and supplemental references.

