## Supplementary material for "Escaping through the predator’s gill cleft: A defensive tactic of juvenile eels after capture by predatory fish": Document S1

**Table and video legends**

Table S1. Summary dataset including the identification and total length of the predator *Odontobutis obscura* (ID_pred and TL_pred, respectively), the total length of the prey *Anguilla japonica* (TL_prey), whether the prey was captured (1: non-captured; 0: captured), whether the captured prey escaped via the gill (1: escaped; 0: swallowed), body part initially attacked (Attack, H: head; B: body; T: tail), time taken from the capture to escape via the gill (Time), and survival for 48 h (Survial_48, 1: survival; 0: death).

Video S1. Video showing the Japanese eel *Anguilla japonica* juvenile escaping through the gill cleft of the predatory fish *Odontobutis obscura*. In this case, the predator did not show apparent resistance behavior, while *A. japonica* escaped though the gill cleft of the predator.

Video S2. Video showing the Japanese eel *Anguilla japonica* juvenile escaping through the gill cleft of the predatory fish *Odontobutis obscura*. In this case, the predator intermittently showed burst movements, while the tail of *A. japonica* was emerging from the gill cleft of the predator.

Video S3. Video in which the Japanese eel *Anguilla japonica* juvenile did not escape after being captured by the predatory fish *Odontobutis obscura*. Note that the video is played at 5×speed.

**Experimental procedures**

We used *A. japonica* juveniles [69.4 ± 8.9 mm (mean ± standard deviation), n = 68] that were bred from recruiting “glass eels” purchased from licensed local fishermen at the Tone River estuary in Chiba, Japan. The *A. japonica* were maintained in glass aquariums (600 × 300 × 360 mm) before starting the experiment and were fed frozen *Chironomus* sp. larvae once every 1–2 days. As predators, we used *O. obscura* (154.0 ± 23.8 mm, n = 4), a common predator in southwestern Japanese rivers, whose distribution range overlaps with that of *A. japonica*. All *O. obscura* samples were collected using a hand net in the Urakami River in Nagasaki, Japan. The predators were kept separately in four glass aquariums (450 × 450 × 450 mm) filled with freshwater to a depth of 130 mm and were fed frozen Manila clams once a day before starting the experiment.

- Experiments were performed in the glass aquariums in which *O. obscura* had been maintained. The water temperature during the experiments was 20.5 ± 0.99 ℃. Because both *A. japonica* and *O. obscura* are nocturnal species [1, 2], the experimental area was covered with black curtains; two infrared lights were used to illuminate the tank, and an infrared video camera (Handycam HDR-XR520; Sony, Tokyo, Japan) was used to record the dorsal view of the fish. One *A. japonica* individual was retrieved from the rearing aquarium using a hand net, quickly weighed after removing surface water with blotting paper, and photographed dorsally to later measure its total length using image analysis software (ImageJ 1.52a, imagej.nih.gov/ij). The photographed individual was then introduced into a PVC pipe (60 mm diameter), set in the center of the experimental aquarium, and acclimated for at least 10 min. After the acclimation period, the trial was started by slowly removing the PVC pipe to release the *A. japonica*, and the predator–prey interaction was recorded using a video camera. If *O. obscura* did not capture *A. japonica* within 20 min, the trial was ended. Four *O. obscura* were used repeatedly, but each *A. japonica* was used only once. We defined “escaped” as a situation in which *A. japonica* that had been captured by the predator (in its buccal cavity) escaped through the gill cleft. When the predator stopped moving its mouth and *A. japonica* did not escape within 3 min afterwards, it was regarded as “swallowed”. When a body part of *A. japonica* (e.g., the tail) emerged from the buccal cavity through the gill cleft, but the individual was finally swallowed by the predator, it was also regarded as “swallowed”. All individuals that escaped through the predator’s gill were kept in a plastic tank (150 × 150 × 150 mm) filled with freshwater to a depth of 70–80 mm for at least 48 h to observe whether they could survive after the experiment.

**Acknowledgements**

We sincerely thank M. Hidaka and S. Tokunaga for their assistance in the collection of *Odontobutis obscura*. This study was funded by Grants-in-Aid for Scientific Research, Japan Society for the Promotion of Science, to Y.K. (19H04936 and 21H02269). Animal care and experimental procedures were approved by the Animal Care and Use Committee of the Faculty of Fisheries, Nagasaki University (Permit No. NF-0046) in accordance with the Guidelines for Animal Experimentation of the Faculty of Fisheries and the Regulations of the Animal Care and Use Committee of Nagasaki University.

**Author contributions**

Conceptualization: Y.H., K.Y., Y.K.; Methodology: Y.H., Y.K.; Software: Y.K.; Formal analysis: Y.H., Y.K.; Investigation: Y.H., Y.K.; Resources: K.Y., Y.K.; Data curation: Y.H., Y.K., Writing - original draft: Y.H., Y.K.; Writing - review & editing: K.Y., Y.K.; Visualization: Y.H., Y.K.; Supervision: Y.K.; Project administration: Y.K.; Funding acquisition: Y.K.
